# Transcriptome analysis in the maize-*Ustilago maydis* interaction identifies maize-line-specific activity of fungal effectors

**DOI:** 10.1101/2020.10.30.361659

**Authors:** S Schurack, JRL Depotter, D Gupta, M Thines, G Doehlemann

**Author notes:** Correspondence to: Gunther Doehlemann.

## Abstract

The biotrophic pathogen *Ustilago maydis* causes smut disease on maize (*Zea mays*) and induces the formation of tumours on all aerial parts of the plant. Unlike in other biotrophic interactions, no gene-for-gene interactions have been identified in the maize-*U. maydis* pathosystem. Thus, maize resistance to *U. maydis* is considered a polygenic, quantitative trait. Here, we study the molecular mechanisms of quantitative disease resistance (QDR) in maize, and how *U. maydis* interferes with its components. Based on quantitative scoring of disease symptoms in 26 maize lines, we performed an RNA-Seq analysis of six *U. maydis*-infected maize lines of highly distinct resistance levels. In accordance with the complex nature of QDR, the different maize lines showed specific responses of diverse cellular processes to *U. maydis* infection. On the pathogen side, our analysis identified 406 *U. maydis* genes being differentially expressed between maize lines, of which 102 encode predicted effector proteins. Based on this analysis, we generated *U. maydis* CRISPR/Cas9 knockout mutants for selected candidate effector sets. Infections of different maize lines with the fungal mutants and subsequent RNA-sequencing identified effectors with quantitative, maize-line-specific virulence functions, and revealed auxin-related processes as a possible target for one of them. Thus, we show that both transcriptional activity and virulence function of fungal effector genes are modified according to the infected maize line, providing new insights into the molecular mechanisms underlying QDR in the maize-*U. maydis* interaction.

## INTRODUCTION

In natural environments, plants are constantly exposed to a variety of potentially pathogenic microbes. To protect themselves, they have evolved multiple layers of immune responses and in turn, microbes have developed so-called effectors to cope with or suppress these immune responses.

Current studies in molecular plant pathology have mainly focused on understanding the molecular mechanisms of qualitative resistance, which is determined by large-effect resistance (R) genes leading to almost complete resistance or susceptibility. This is often conferred by nucleotide-binding leucine-rich repeat (NLR) receptors (Dangl and Jones 2001; McHale et al. 2006; Cui et al. 2015). In contrast, quantitative disease resistance (QDR) still remains poorly understood (Poland et al. 2009; Roux et al. 2014; Corwin and Kliebenstein 2017), even though it determines the outcome of the majority of plant-pathogen interactions in crops and natural populations (Bartoli and Roux 2017). In QDR, many genes with small to moderate effects lead to a continuous distribution of susceptible to resistant phenotypes (Poland et al. 2009; St.Clair 2010; Roux et al. 2014; Niks et al. 2015).

Many QDR loci have been mapped in the past, but the underlying complex genetic architecture has limited the molecular characterisation of mechanisms involved (Corwin and Kliebenstein 2017). Still, several QDR genes with various functions have been cloned recently. In several cases, kinases have been shown to play important roles in QDR. Two maize wall-associated kinases, ZmWAK-RLK1 and ZmWAK, confer QDR to northern corn leaf blight and head smut, respectively (Hurni et al. 2015; Zuo et al. 2015). Other QDR genes encode putative transporters, the ABC (adenosine triphosphate [ATP]-binding cassette) transporter encoded by Lr34 confers resistance to diverse fungal pathogens in wheat (Krattinger et al. 2009). A caffeoyl-CoA O-methyltransferase connected to lignin production was shown to confer QDR to various necrotrophic maize pathogens (Yang et al. 2017). Other genes identified in QDR correspond to previously unidentified defence genes, such as the soybean wound-inducible domain protein WI12, the soybean serine hydroxymethyltransferase RHG4 and the rice proline-containing protein Pi21 (Fukuoka et al. 2009; Cook et al. 2012). In rare cases, NLR genes can also underlie QDR (Poland et al. 2009; Barbacci et al. 2020), which led to the hypothesis that allelic variants, i.e. weak alleles, of R genes can cause incomplete resistance. Thus, compared to qualitative resistance, the molecular functions underlying QDR seem to be highly diverse. In general, QDR can be hypothesised as an intricate network integrating various response pathways to pathogen determinants and environmental factors (Roux et al. 2014). Additionally, studies of QDR by RNA-Seq approaches have indicated highly interconnected and multifaceted defence responses, which were mostly distinct from functions previously identified for plant immunity (Kebede et al. 2018; Pan et al. 2018; Delplace et al. 2020).

The biotrophic fungus *Ustilago maydis* causes smut disease in maize (*Zea mays*). Characteristic disease symptoms are tumours that can be formed on all above-ground organs of maize plants in less than two weeks (Basse and Steinberg 2004; Kämper et al. 2006). *U. maydis* has advanced to a model for biotrophic plant pathogens due to its rapid symptom development, very compact genome, easy *in-vitro* cultivation and accessibility to genetic manipulation (Kämper 2004; Brefort et al. 2009; Dean et al. 2012; Schuster et al. 2016; Zuo et al. 2019; Zuo et al. 2020). The infection cycle of *U. maydis* is initiated by recognition and fusion of sporidia with compatible mating types, leading to a morphological switch from yeast-like haploid cells to diploid filaments (Bölker et al. 1992; Spellig et al. 1994). The generation of the solopathogenic strain SG200, derived from a field isolate collected in Minnesota, USA, and genetically engineered to be able to form infectious filaments without prior mating, has greatly facilitated the investigation of *U. maydis* pathogenic development (Kämper et al. 2006). The genome of *U. maydis* is predicted to encode 553 secreted effector proteins, of which the majority lacks known functional or structural domains (Dutheil et al. 2016). Many effectors reside in clusters in the genome, are expressed specifically during biotrophic development compared to axenic culture, and contribute to virulence (Kämper et al. 2006; Müller et al. 2008; Skibbe et al. 2010; Schirawski et al. 2010; Schilling et al. 2014). However, so far only a few individual effectors with large effects on virulence have been functionally characterised (Doehlemann et al. 2009; Djamei et al. 2011; Tanaka et al. 2014; Redkar et al. 2015; Ma et al. 2018; Ökmen et al. 2018, Sharma et al. 2019). Despite *U. maydis* being the predominant model organism of biotrophic plant pathogens, resistance to *U. maydis* has been rarely described (Lübberstedt et al. 1998; Baumgarten et al. 2007). Unlike in other biotrophic interactions, no gene-for-gene interactions have been identified in the *U. maydis-maize* pathosystem. Crosses of *U. maydis*-resistant and -susceptible maize lines have previously indicated that *U. maydis* resistance is a polygenic, quantitative trait (Immer 1927; Hoover 1932). Several QDR loci that contribute to *U. maydis* infection frequency and severity, have been identified and some studies have suggested that specific loci may contribute to *U. maydis* resistance in an organ-specific manner (Lübberstedt et al. 1998; Baumgarten et al. 2007). Interestingly, several QDR loci conferring resistance to *U. maydis* contain genes with a known role in defence against pathogens, such as NLRs, a pathogenesis-related protein, a chitinase, a basal antifungal protein, and a wound-inducible protein (Baumgarten et al. 2007; Brefort et al. 2009). For one *U. maydis* effector, ApB73, a maize line-specific virulence function has been observed (Stirnberg and Djamei 2016). This suggests that the fungus’ effectors might target certain QTL gene products. However, the molecular basis of QDR in maize and how *U. maydis* interferes with its components is still mostly unknown.

Extensive transcriptome analyses have revealed organ-, cell type-as well as stage-specific expression patterns of the *U. maydis* effector gene repertoire (Skibbe et al. 2010; Lanver et al. 2018; Matei et al. 2018). Despite these efforts, how gene expression is influenced by host lines of quantitatively different resistance levels remains to be elucidated. Such knowledge would help to draw a more comprehensive picture of *U. maydis* virulence.

In this study, we analysed the transcriptome of *U. maydis* infecting six maize lines of quantitatively differing resistance levels via RNA-Seq. This offered unprecedented insights into transcriptional changes associated with host disease resistance. Within the 406 *U. maydis* genes identified to be expressed in a host genotype-dependent manner, effectors were significantly enriched, suggesting their predominant role in response to quantitatively different host resistance levels. On the plant side, in line with the complex nature of QDR, we found genes of various functional classes to be associated to QDR against *U. maydis*

## RESULTS

### *U. maydis* disease development in different maize lines

To investigate quantitative disease resistance in the maize-*U. maydis* interaction, we first evaluated the susceptibility of different maize lines to *U. maydis* infection. For this, *U. maydis* resistance levels were assessed in the 26 inbred founder lines of the Nested Association Mapping recombinant inbred lines (NAM RILs) (Yu et al. 2008; McMullen et al. 2009), a set of maize lines selected to represent world-wide maize diversity. In addition, the sweet corn Early Golden Bantam (EGB) was used, which is mostly used in *U. maydis* research (Zuo et al. 2019). Seedling infections were performed in three independent biological replicates under controlled conditions with an average of 102 plants per line being scored for *U. maydis* disease symptoms (Fig. 1A). In this experimental set-up, resistance levels were highly diverse and ranged from very susceptible to very resistant (>94% vs. <35% tumours, respectively), while no maize line showed complete resistance to *U. maydis* infection (Fig. 1A). Agglomerative hierarchical clustering of disease indices as a measure of *U. maydis* infection severity identified five susceptibility groups (Fig. 1B). Two groups consisted only of the most resistant line CML322 and of the most susceptible line Tx303, respectively, and three groups were of comparable sizes, indicating a mostly even distribution of *U. maydis* resistance levels within the NAM founder lines and EGB. The *U. maydis* SG200 strain used in this study was derived from a field isolate from a temperate region (Minnesota, USA; Kämper et al. 2006). Strikingly, among the maize lines with highest susceptibility, most were local to regions close to the origin of SG200 (e.g. Oh43 from Ohio, Mo18w from Missouri, Il14H from Illinois). In contrast, all four most resistant maize lines were of tropical origin (CML322, NC350, NC358, Ki3). Thus, maize lines of close provenance to SG200 were generally more susceptible, indicating a possible adaptation of the local *U. maydis* strain to the local host lines. From each group, 1-2 lines were chosen based on resistance level, provenance, growth soundness and seed production for subsequent investigations (CML322, B73, EGB, Ky21, Oh43 and Tx303).

**Figure 1.**
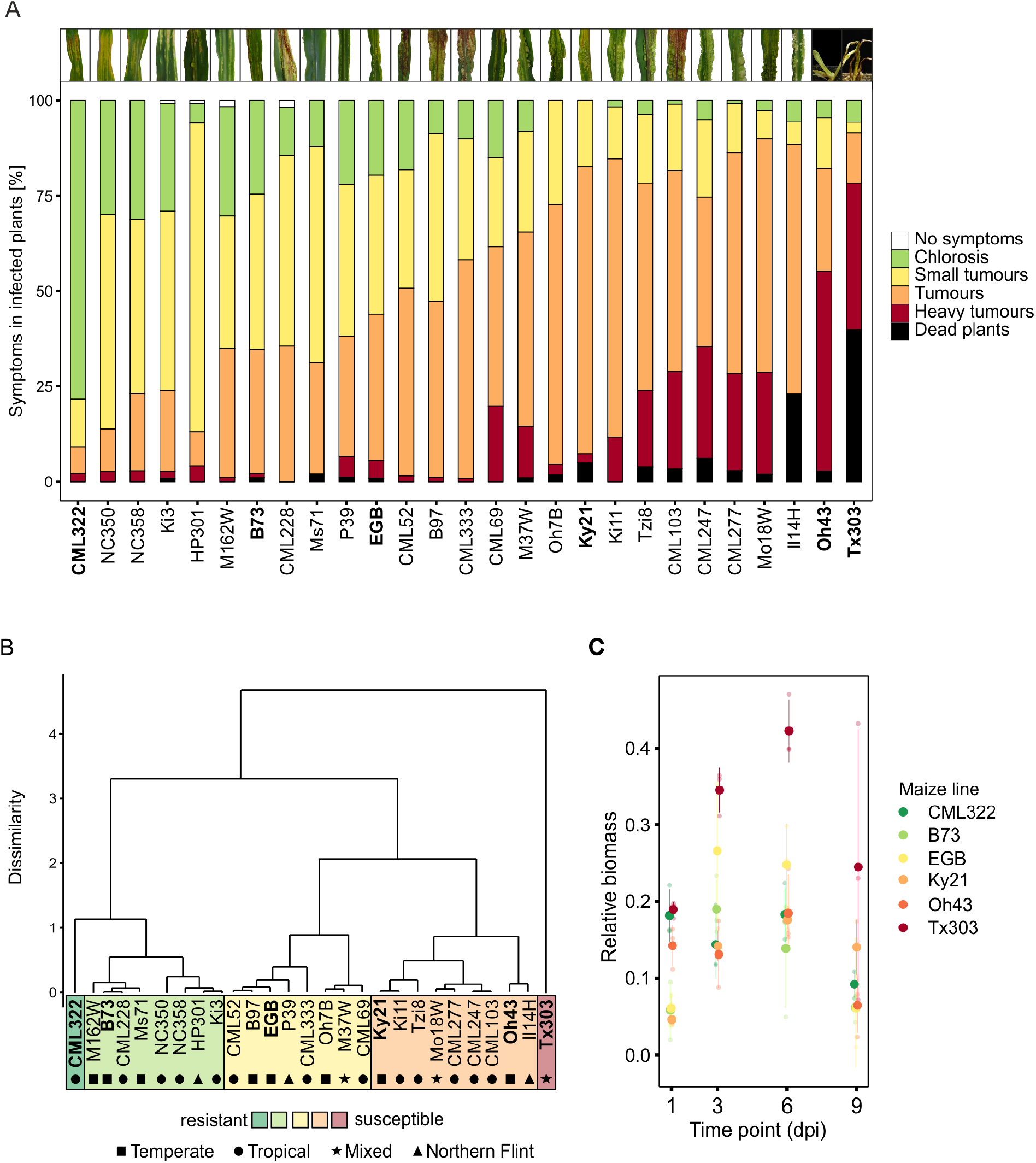
*U. maydis* disease development in the 26 maize NAM founder lines and EGB. **A) Disease symptom classification.** Maize seedlings were infected with *U. maydis* SG200 at the three-leaf stage. Three independent experiments were performed and the average values are expressed as percentage of the total number of infected plants. Disease symptom classification was done 12 days post infection (dpi) as described in Redkar & Doehlemann (2016a). Average number of infected plants per line: 102. Maize lines selected for RNA-Seq are highlighted in bold. Representative pictures of infected leaves at 12 dpi for each maize line are given at the top. **B) Agglomerative hierarchical clustering of disease indices.** Clustering is based on Euclidean distances of disease indices using complete linkage clustering. Maize lines selected for RNA-Seq are highlighted in bold. The maize lines’ provenances are depicted by black symbols. **C) Fungal biomass quantification based on the amount of genomic DNA.** A qPCR with plant-specific (GAPDH) and fungus-specific (ppi) primers was performed at 1, 3, 6 and 9 dpi in the maize lines selected for RNA-Seq. Solid points give mean ratios of fungal DNA to plant DNA (2^−ΔCt^) of three biological replicates, transparent points give individual values, error bars denote the standard deviation.

To further characterise disease progression of *U. maydis* within the different maize lines and to select a time point suitable for transcriptome analysis, we assessed relative fungal biomass by qPCR using gDNA (Fig. 1C) and visualised fungal growth within leaf tissue by WGA-AF488/propidium iodide co-staining throughout the infection process at 1, 3, 6 and 9 days post infection (dpi) (Supplementary Fig. S1).

At 1 and 3 dpi, relative fungal biomass did not differ significantly between the maize lines. At 6 dpi however, fungal biomass in Tx303, the most susceptible maize line, was increased approximately two-fold compared to the other maize lines. In line with previous observations, relative fungal biomass decreased at the late infection time point (9 dpi), which might be due to an impaired teliospore formation in the genetically engineered haploid SG200 strain (Lanver et al. 2018).

At the microscopic level, the infection progress was comparable for 1 and 3 dpi in all maize lines as well. At 6 dpi, strong differences could be observed, as for CML322, the most resistant maize line, hyphae were still only mostly proliferating, whereas for the more susceptible maize lines, fungal aggregates, fragmented hyphae and enlarged maize cells were visible. Size and number of fungal aggregates and maize cell enlargements increased with susceptibility levels of the maize lines. Based on these fungal quantification and microscopic growth data, the 3 dpi time point was chosen for transcriptome analysis. At this time point, the different maize lines showed comparable growth of biotrophic hyphae while levels of fungal colonisation allowed sufficient coverage of *U. maydis* genes by RNA sequencing.

### Transcriptome analysis of *U. maydis* infecting maize lines of distinct disease resistance levels

To analyse the gene expression changes induced by different host genotypes of distinct resistance levels, maize seedlings of CML322, B73, EGB, Ky21, Oh43 and Tx303 were infected with *U. maydis* SG200 and water (mock control). Infected and mock-treated leaf sections were collected 3 dpi in biological triplicates and their transcriptome was subsequently analysed via RNA-Seq. After filtering for low expression, 6284 of 6766 *U. maydis* genes remained for the analysis and were considered to be expressed in our samples (93%). To evaluate variability between the samples, we made a multidimensional scaling (MDS) plot (Fig. 2A). To additionally examine whether the infection stage in the different maize lines was comparable and to demonstrate that gene expression differences were not caused by faster infection progression in the more susceptible maize lines, we included transcriptome data previously published by Lanver et al. (2018), where the maize line EGB was infected with the more virulent *U. maydis* wild type crossing FB1xFB2 and the fungal transcriptome was mapped during different stages of the infection process. All our samples clustered with the 2 dpi samples of Lanver et al. (2018), which likely reflects the slower disease progression of SG200 compared to FB1xFB2. Again, this showed no pronounced differences in infection progression between the different maize lines at the time point tested.

**Figure 2.**
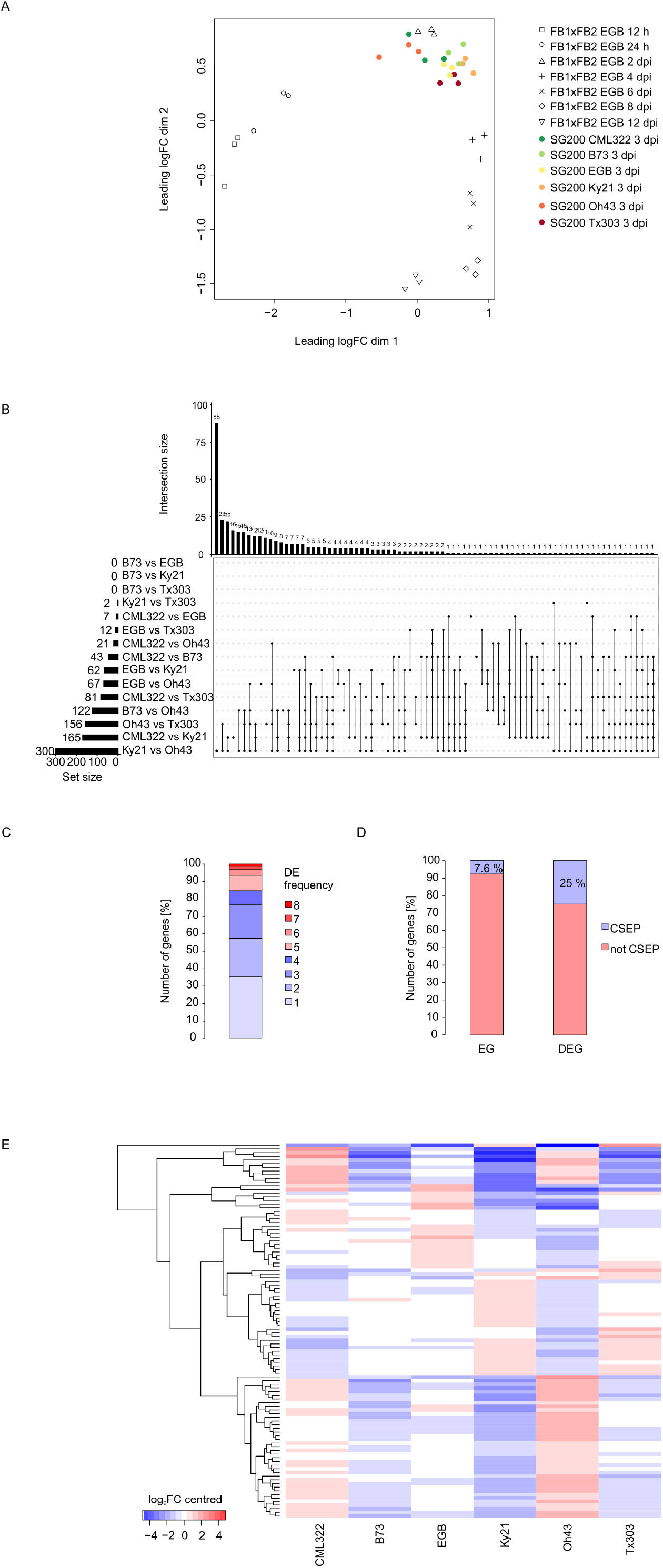
RNA-Seq analysis of *U. maydis* infecting maize lines of differing disease resistances. **A) MDS plot of *U. maydis* RNA-Seq data.** The top 1000 variable genes were used to calculate pairwise distances between the samples. FB1xFB2 RNA-Seq data were previously published and represent different time points in the *U. maydis* disease cycle in EGB (Lanver et al. 2018). MDS: multidimensional scaling. **B) UpSet plot of the distribution of differentially expressed *U. maydis* genes across maize lines.** Genes with a log_2_ expression fold change >0.5 and adjusted p value <0.05 were considered differentially expressed (DE). In total, 406 of 6284 expressed genes were differentially expressed between maize genotypes. Number of DE genes (DEGs) for each of the 15 possible comparisons is given by set size (horizontal bars). Overlaps of DEGs between comparisons are depicted by connected black dots. Size of overlaps is given by intersection size (vertical bars). **C) Number of differentially expressed genes by frequency of differential expression within comparisons.** The categories of the bar plot give the percentage of all DEGs that are DE in the indicated number of comparisons. DE: differential expression. **D) Enrichment of CSEPs in differentially expressed genes.** Portion of candidate secreted effector proteins (CSEPs) in all 6284 expressed *U. maydis* genes compared to the portion of CSEPs in genes DE between maize genotypes. Within DE genes, CSEPs show a 3.3-fold enrichment (hypergeometric test, p value 5.65e-30). EG: expressed genes. DEG: differentially expressed genes. CSEP: candidate secreted effector protein. **E) Expression profile of differentially expressed *U. maydis* CSEPs across maize lines.** Heatmap shows log_2_ expression fold changes compared to mean expression across all samples. CSEP: candidate secreted effector protein.

To analyse whether *U. maydis* gene expression is influenced by the colonised maize line, we compared expression in all 15 possible pairs of the six different maize lines. This analysis showed that in total 406 of the 6284 expressed genes (6.4%) were differentially expressed (log_2_ expression fold change >0.5, adjusted p value <0.05, Supplementary Data Set S1) in at least one of the 15 comparisons. The number of differentially expressed *U. maydis* genes (DEGs) ranged from zero to 300 genes in the different comparisons and only a few genes were differentially expressed in several of the 15 comparisons (Fig. 2B, C). The majority of DEGs was only differentially expressed in one to three comparisons (approx. 75%) and only 1% of DEGs was differentially expressed in more than half of the comparisons, which suggests that not a shared set of genes is responsive to different host environments, but that different maize lines lead to rather diverse gene expression changes. Strikingly, amongst the 406 DEGs, 102 encode candidate secreted effector proteins (CSEPs, Dutheil et al. 2016), which represents a significant 3.3-fold enrichment (hypergeometric p value 5.65e-30) (Fig. 2D). A heatmap based on the expression profiles of the 102 line-specific CSEPs shows distinct groups of CSEPs with similar expression patterns (Fig. 2E, Supplementary Data Set S2). Of the 102 CSEPs, one group of 38 genes is upregulated on the most resistant maize line CML322 and downregulated in more susceptible maize lines, except for Oh43, while another group of 29 CSEPs shows an opposite expression pattern. Besides these two main expression patterns, some CSEPs show no clear correlation to the resistance level. Consequently, a dominant expression pattern that underlies all maize line-specific CSEPs cannot be observed.

### Weighted gene co-expression analysis (WGCNA) of *U. maydis* genes during infection of maize lines of distinct disease resistance levels

To elucidate which processes could be involved in colonising maize lines of distinct resistance levels, we assessed for which *U. maydis* genes expression correlated to the resistance level of the colonised maize line. To this end, we first performed a weighted gene co-expression network analysis (WGCNA), using the *U. maydis* expression data of the different maize lines. WGCNA identifies modules of co-expressed genes and represents the modules by summary expression profiles, referred to as the module eigengene (Zhang and Horvath 2005; Langfelder and Horvath 2008). This analysis identified 11 colour-coded modules with differential expression profiles of the module eigengenes, ranging in size from 1073 (‘turquoise’) to 65 genes (‘purple’) (Fig. 3A, Supplementary Data Set S3). Subsequently, in order to identify modules associated with the severity of the infection, the correlation of each module eigengene with the disease indices of the different maize lines was calculated (referred to as gene significance, GS) (Fig. 3B). The ‘purple’ module showed a significant positive correlation (GS >0.5, p value <0.05) and the ‘greenyellow’ module showed significant negative correlation to the disease index (GS <-0.5, p value <0.05), i.e. expression of genes in the ‘purple’ module is higher in more susceptible maize lines and expression of genes in the ‘greenyellow’ module is higher in more resistant maize lines. To evaluate which biological processes were associated with the colonisation of more resistant and more susceptible maize lines, the ‘purple’ and ‘greenyellow’ modules were subjected to enrichment analysis of Gene Ontology (GO) terms and CSEPs (Ashburner et al. 2000; The Gene Ontology Consortium 2017), (Fig. 3C, Supplementary Data Sets S4,S5). In summary, mostly ion transport processes were significantly enriched in the ‘purple’ module. Ion transmembrane transport through H^+^-ATPases is a crucial driving force for nutrient exchange between host plants and fungi (Palmgren 1990; Gianinazzi-Pearson et al. 1991; Sondergaard et al. 2004; Wang et al. 2014). Furthermore, different nutrient transporters were found to be important virulence factors tied to biotrophic development in *U. maydis* (Lanver et al. 2018). As indicated by the enrichment of ion transport processes in the module with higher gene expression in more resistant maize lines, different availability of nutrients in more resistant vs. more susceptible maize lines could therefore be involved in QDR to *U. maydis*. Additionally, ‘oxidation-reduction’ was the GO term with the most genes. Oxidation-reduction processes are involved in metabolism as well, but can also have a signalling function or be related to detoxification of reactive oxygen species (ROS). In the ‘greenyellow’ module, all significantly enriched GO terms were related to carbohydrate metabolism. In addition, CSEPs were significantly enriched in this module and represented the biggest category. Carbohydrate utilization has been directly linked to plant cell wall degradation in other plant pathogenic fungi (Tonukari et al. 2000; Ospina-Giraldo et al. 2003). Since carbohydrate metabolism is enriched in the module with higher gene expression in more resistant maize lines, it could be speculated if the fungus might need to overcome enhanced cell wall reinforcements as part of increased resistance. The enrichment of CSEPs in this module might represent an attempt of the fungus to suppress enhanced defence mechanisms in more resistant host lines.

**Figure 3.**
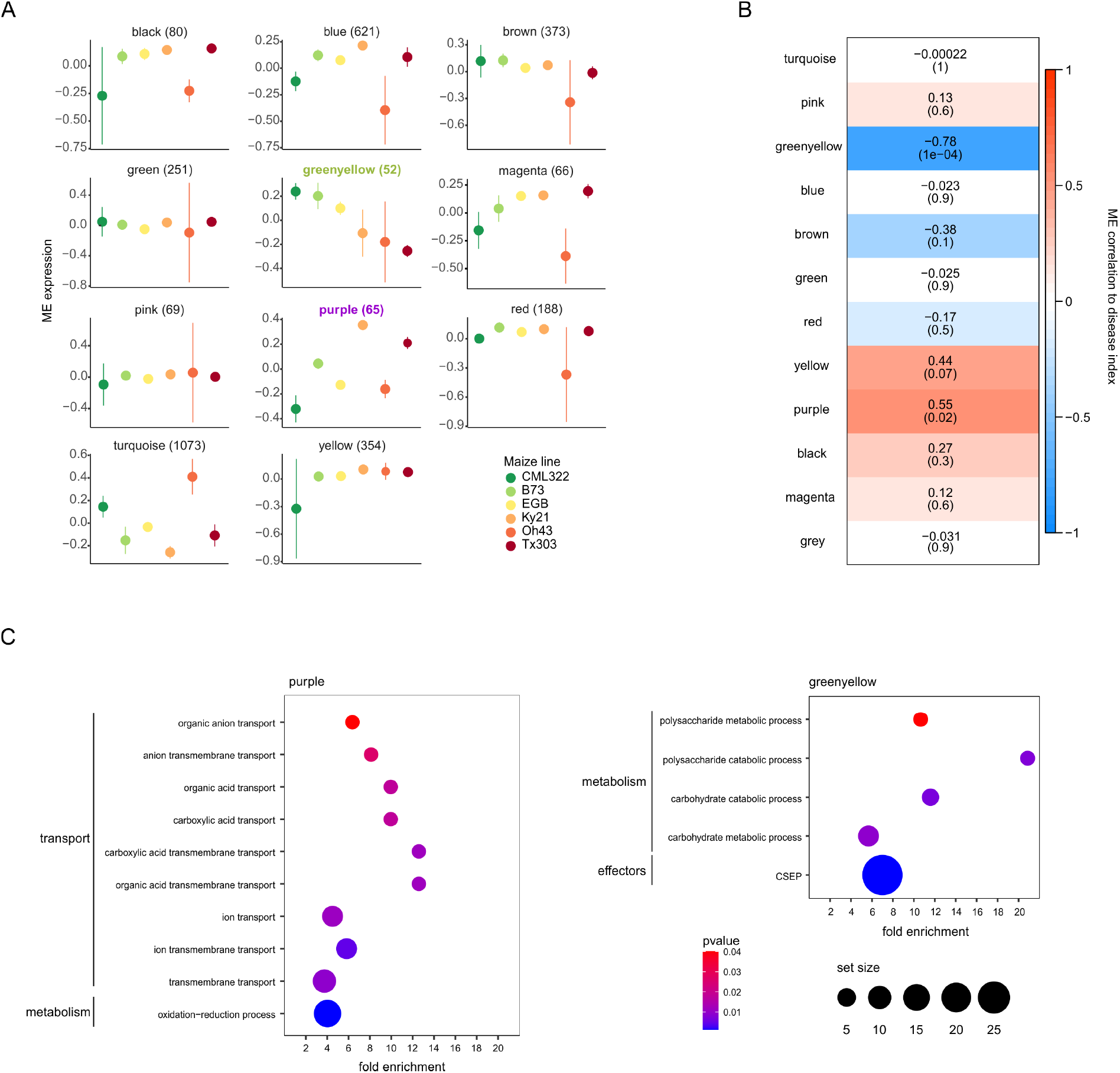
Weighted gene co-expression analysis (WGCNA) of *U. maydis* genes during infection of maize lines of differing disease resistances. **A) Modules of co-expressed *U. maydis* genes across maize lines.** The RNA-Seq data was subjected to WGCNA to detect modules of co-expressed genes. Each plot represents the expression profile of the module eigengene, which can be considered as representative of the expression of the respective co-expression module. Error bars indicate standard deviation of three biological replicates. The modules are named according to their colour, and the size of each module is given in parentheses. Modules significantly correlated with disease index are highlighted in bold and their respective colour. **B) Module-disease index association**. Correlation of modules of co-expressed genes with the disease index of the colonised maize line. Numbers in the heatmap show the correlations with disease index and p values in parentheses for the respective module eigengene (ME). Correlation was considered significant for correlation >0.5 or <-0.5 and p value <0.05. **C) GO and CSEP enrichments in modules correlated with disease index.** GO biological process terms and additionally CSEPs (candidate secreted effector proteins) were tested for significant enrichment in the purple (positive correlation to disease index) and greenyellow (negative correlation to disease index) modules. Gene sets were considered significantly enriched for p <0.05 (hypergeometric test). Dot size is representative of the number of analysed genes in the respective term. Only genes with a gene significance to disease index of >0.5 (purple) or <-0.5 (greenyellow) and p value <0.05 were considered for the analysis and only terms with a set size ≥2 are shown.

### Transcriptome analysis of *U. maydis*-infected maize lines of distinct disease resistance levels

In order to identify maize genes involved in QDR to *U. maydis*, we analysed genotype-dependent transcriptional changes in response to *U. maydis* via RNA-Seq. Of all 63477 maize annotated loci, 40056 were expressed in our samples (63%). To assess the variability between the samples we used a multidimensional scaling plot (Fig. 4A). *U. maydis*-infected and control samples formed two distinct groups, within which the samples of each maize line clustered together, indicating both treatment-specific and genotype-specific expression patterns. To identify genes which differentially respond to *U. maydis* infection between maize lines, we compared expression fold changes of the *U. maydis-infected* samples to the respective control samples for all 15 possible pairs of six different maize lines (i.e. difference between genotypes in response to infection). This analysis showed that in total 8675 of 40056 transcripts (22%) responded differentially to *U. maydis* infection (log_2_ expression fold change >0.5, adjusted p value <0.05, Supplementary Data Set S6) in at least one of the 15 comparisons. The number of DEGs ranged from 358 to 1283 genes in the different comparisons and the fraction of genes differentially expressed in several of the 15 comparisons was very small (Fig. 4B). Around 50% of DEGs were differentially expressed in only one of the comparisons and only 4% of DEGs were differentially expressed in more than half of the comparisons. Together, this shows that genes differentially responding to *U. maydis* infection are highly diverse between maize lines.

**Figure 4.**
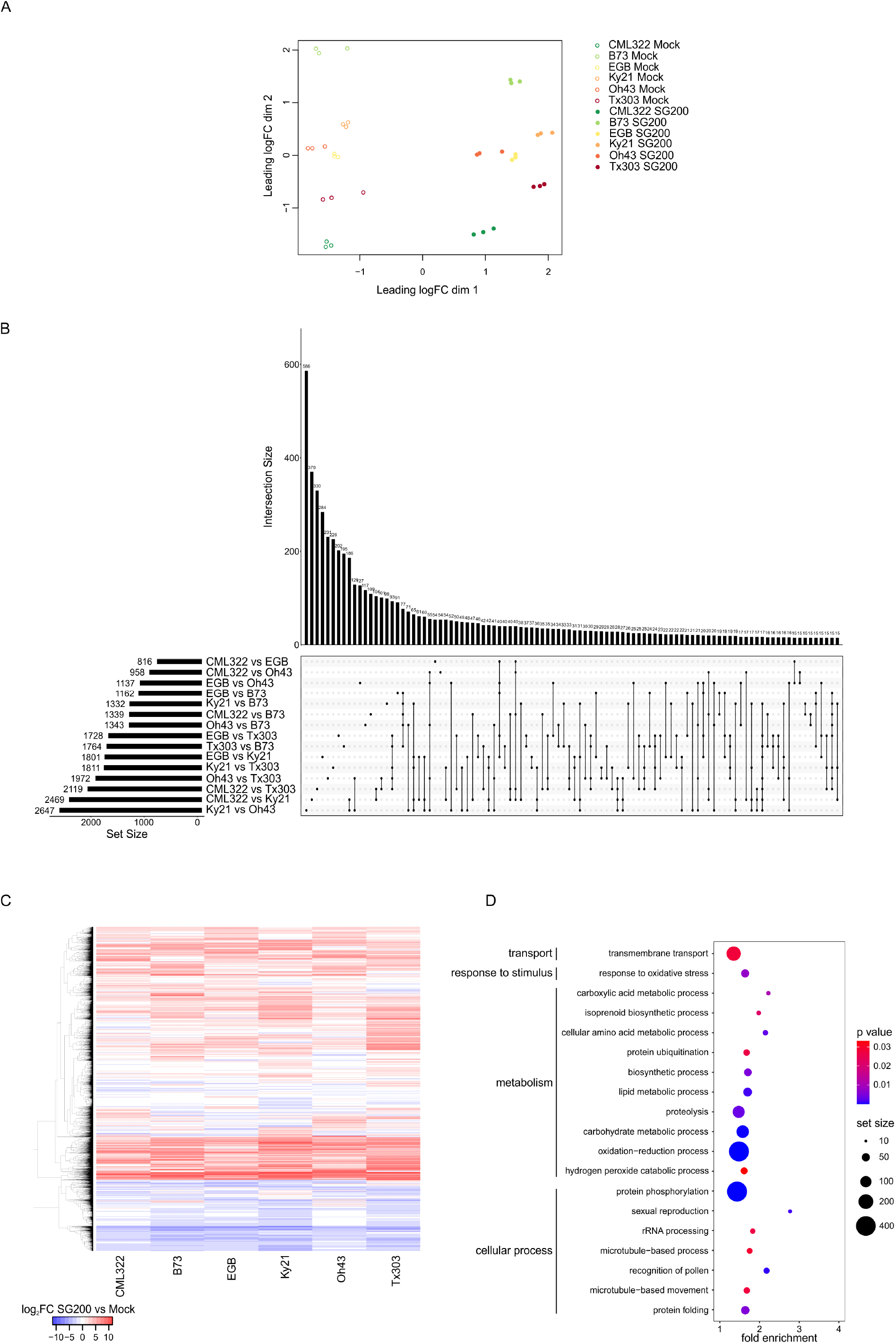
RNA-Seq analysis of maize gene expression in response to *U. maydis*. **A) MDS plot of maize RNA-Seq data.** The top 5000 variable genes were used to calculate pairwise distances between the samples. MDS: multidimensional scaling. **B) UpSet plot of the distribution of genes differentially expressed between maize lines in response to *U. maydis*.** Genes with a log_2_ expression fold change >0.5 of the gene expression changes and adjusted p value <0.05 were considered differentially expressed (difference between genotypes in response to infection). In total, 8675 of 40056 expressed genes were differentially responding to *U. maydis* between maize genotypes. Number of differentially expressed genes (DEGs) for each of the 15 possible comparisons is given by set size (horizontal bars). Overlaps of DEGs between comparisons are depicted by connected black dots. Size of overlaps is given by intersection size (vertical bars). **C) Expression profile of differentially expressed maize genes in response to *U. maydis*.** Heatmap shows log_2_ expression fold changes of SG200-infected vs mock-treated samples. **D) GO enrichments in differentially expressed maize genes.** GO biological process terms were tested for significant enrichment in all genes differentially expressed between maize lines in response to *U. maydis*. Gene sets were considered significantly enriched for p <0.05 (hypergeometric test). Dot size is representative of the number of analysed genes in the respective term. Only terms with a set size ≥10 are shown.

To identify biological processes which were associated with the maize line-specific gene expression responses, all DEGs were subjected to enrichment analysis of GO terms (Ashburner et al. 2000; The Gene Ontology Consortium 2017), highlighting processes involved in transport, response to stimulus, cellular processes and metabolism (Fig. 4D, Supplementary Data Set S8). The GO terms with most genes were ‘transmembrane transport’ as well as ‘oxidation-reduction’ and ‘protein phosphorylation’, which could indicate a special importance of these processes in genes differentially regulated in response to *U. maydis* between maize lines. Transport processes play a pivotal role in signalling, nutrient uptake as well as growth and development. Oxidation-reduction processes are involved in metabolism but can also have a signalling function. Protein phosphorylation occurs during kinase signalling processes. A predominant role of genes related to metabolism as well as kinase-signalling cascades for QDR has been proposed before (Delplace et al. 2020). Taken together, this suggests that maize line-specific responses to *U. maydis* involved various cellular activities, consistent with the complex nature of QDR. To examine if the maize DEGs include genes associated with other forms of immunity, we compared *Arabidopsis thaliana* orthologues of the DEGs with *A. thaliana* genes previously found to be linked to pathogen-associated molecular pattern (PAMP)-triggered immunity (PTI) and/or effector-triggered immunity (ETI) responses (Dong et al. 2015; Hatsugai et al. 2017; Mine et al. 2018). Of the 3264 DEG *A. thaliana* orthologues, only about 11% (363 and 360 genes) were found in common with either ETI- and/or PTI-associated genes (Supplementary Fig. S2A). This result might suggest that processes differentially regulated between maize lines in response to *U. maydis* are likely distinct from canonical ETI and PTI pathways.

To assess which processes could be connected to *U. maydis* resistance or susceptibility, the correlation of expression changes between *U. maydis*-infected and mock of each gene of the respective maize line to the disease index was calculated (Supplementary Data Set S7). All DEGs were then filtered for genes with a significant positive (GS >0.5 and p value <0.05, Fig. 5A,) or negative (GS <-0.5 and p value <0.05, Fig. 5B) correlation to the disease index. Next, these two sets of genes were again subjected to enrichment analysis of GO terms (Fig. 5A, B, Supplementary Data Sets S9, S10). In the DEGs with positive correlation to the disease index, i.e. upregulated in more susceptible maize lines, enrichments were found in four main cellular activities: cellular processes, response to stimulus, transport, and metabolism (Fig. 5A). The enriched GO term with the largest number of genes was ‘protein phosphorylation’, one of the most important cellular regulatory mechanisms involved in signal transduction. Furthermore, biological process terms that can be linked to cell division processes (‘DNA replication’, ‘microtubule-based movement’) and ‘sexual reproduction’/’recognition of pollen’ were significantly enriched. In DEGs negatively correlated to the disease index, i.e. genes upregulated in the more resistant maize lines, enrichments were found in transport and metabolism (Fig. 5B). The enriched GO term with the largest number of genes was ‘translation’, and a process that could be involved in photosynthesis (‘porphyrin-containing compound biosynthetic process’) was most strongly enriched.

**Figure 5.**
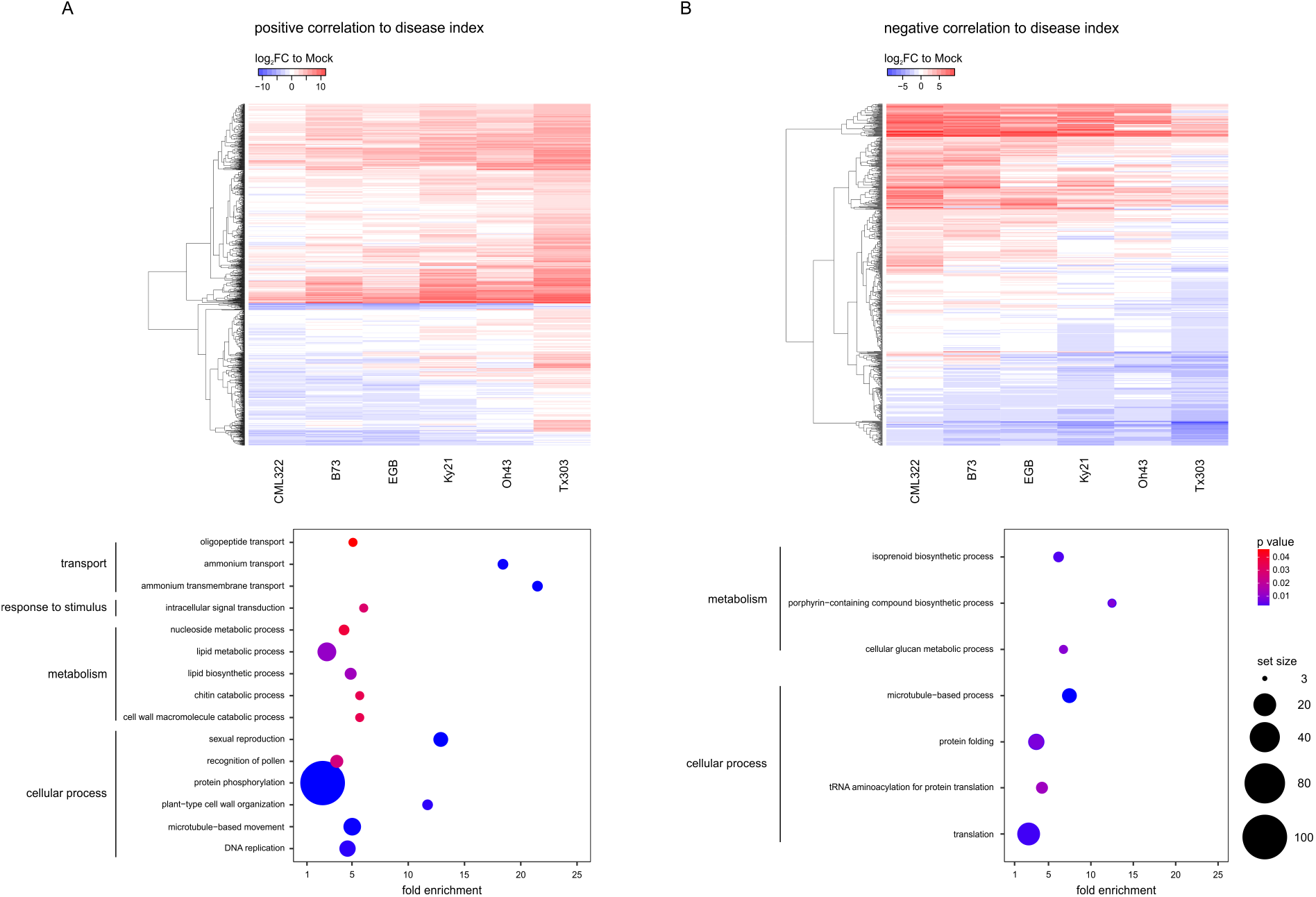
Correlation of maize gene expression to *U. maydis* resistance levels. **A) Expression profile and enrichments of genes positively correlated with the disease index**. Genes with a gene significance for the disease index >0.5 and p <0.05 were considered to be significantly positively correlated to the disease index. Heatmap shows log_2_ expression fold changes of SG200-infected vs mock-treated samples. GO biological process terms were tested for significant enrichment in all genes differentially expressed between maize lines in response to *U. maydis* and positively correlated to the disease index. Gene sets were considered significantly enriched for p <0.05 (hypergeometric test). Dot size is representative of the number of analysed genes in the respective term. Only terms with a set size ≥3 are shown. **B) Expression profile and enrichments of genes negatively correlated with the disease index**. Genes with a gene significance for the disease index <−0.5 and p <0.05 were considered to be significantly negatively correlated to the disease index. Heatmap shows log_2_ expression fold changes of SG200-infected vs mock-treated samples. GO biological process terms were tested for significant enrichment in all genes differentially expressed between maize lines in response to *U. maydis* and negatively correlated to the disease index. Gene sets were considered significantly enriched for p <0.05 (hypergeometric test). Dot size is representative of the number of analysed genes in the respective term. Only terms with a set size ≥3 are shown.

The reactivation of cell division processes including DNA replication are crucial for formation of *U. maydis-induced* tumours (Doehlemann et al. 2008b; Redkar et al. 2015; Matei et al. 2018; Villajuana-Bonequi et al. 2019). Hence, enrichment of such processes in more susceptible maize lines is not surprising since there, *U. maydis* induces more and larger tumours compared to the more resistant lines. Suppression of photosynthesis-associated genes is a typical process in *U. maydis*-infected tissue, where the normal development from sink to source is prevented (Doehlemann et al. 2008a). The finding that processes related to photosynthesis were enriched within maize genes upregulated in more resistant maize lines indicates that here, the induction of such genes is less reduced by *U. maydis* infection.

### Identification of *U. maydis* CSEPs targeting components of QDR

As *U. maydis* genes encoding CSEPs were enriched both in genes differentially expressed between maize lines, as well as in the co-expression module correlated to infection severity, we decided to investigate whether line-specifically expressed CSEPs also have line-specific functions for virulence. To this end, 12 candidate maize line-specific (Mls) CSEP genes were selected from all 102 differentially expressed CSEPs based on a high log_2_ expression fold change and an expression pattern with higher expression in resistant and lower in susceptible maize lines or vice versa (sum of log_2_ expression fold change across all samples >2; Fig. 6A). CSEPs with similar expression patterns were targeted for simultaneous knock-out in the SG200 background using the CRISPR/Cas9 system (Fig. 6A, B, Schuster et al., 2016). Plant infections with the generated *U. maydis* mutant strains identified line-specific virulence functions for the CSEP genes UMAG_02297 and/or UMAG_05027 (Fig. 6C). While virulence of the double mutant KO_UMAG_02297/KO_UMAG_05027 was not reduced on B73 or EGB, a significant reduction was observed on CML322 and Oh43. For reasons of seed availability, subsequent analyses of the mutants were focussed on the maize line CML322. Here, the virulence defect could be restored by introducing single copies of both genes into the *ip* locus of the double mutant strain, demonstrating specificity of the observed virulence reduction (Fig. 6C).

**Figure 6.**
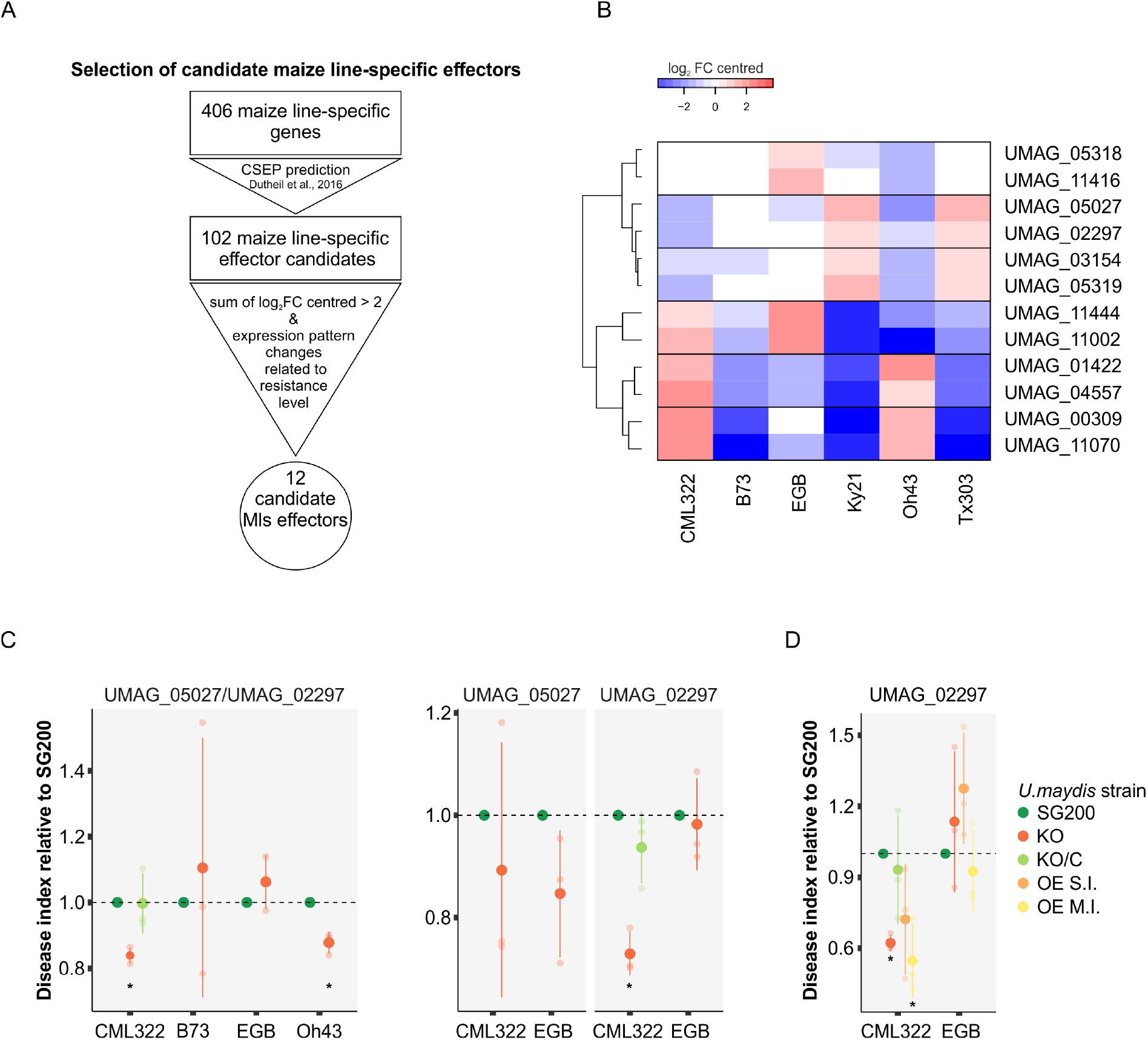
Expression pattern and virulence function of candidate maize line-specific effectors. **A) Selection of maize line-specific effector candidates for functional characterization. B) Expression profile of selected maize line-specific effector candidates across maize lines.**Heatmap shows log_2_ expression fold changes compared to mean expression across all samples. **C) Virulence function of candidate maize line-specific effectors.** Double and single knock-out (KO) mutant strains of selected maize line-specific effectors were injected into maize seedlings of the indicated line and symptoms were scored 12 days post infection (dpi). Gene names are given at the top. KO refers to the respective CRISPR/Cas9 knock-out strain. Gene names separated by slash indicate double KO of these genes. KO/C indicates that a single copy of the respective gene(s) was introduced into the KO strain for complementation. Disease indices reflect disease symptom severity and are shown in relation to SG200, which was set to unity. Asterisks label significant reduction in disease index compared to SG200 (student’s t-test, p<0.05). All experiments were performed in three independent biological replicates. Average number of infected plants per strain and maize line: 89. **D) Impact of UMAG_02297 overexpression on virulence.** SG200, KO_UMAG_02297, KO_UMAG_02297/C and OE_UMAG_02297 strains were injected into CML322 and EGB seedlings and symptoms were scored 12 dpi. OE: overexpression. S.I.: single integration. M.I.: multiple integration. Disease indices reflect disease symptom severity and are shown in relation to SG200, which was set to unity. Asterisks label significant reduction in disease index compared to SG200 (student’s t-test, p<0.05). All experiments were performed in three independent biological replicates. Average number of infected plants per strain and maize line: 86.

In order to assess if both or only one of the genes contribute to maize line-specific virulence of KO_UMAG_05027/KO_UMAG_02297 on CML322, single KO mutants of UMAG_02297 and UMAG_05027 were tested for virulence on EGB and CML322. This experiment showed that UMAG_02297 alone, but not UMAG_05027, was necessary for full virulence on CML322. The virulence defect of KO_UMAG_02297 could be restored by introducing a single copy of UMAG_02297 into the *ip* locus of the mutant strain (Fig. 6C). In addition, a line-specific virulence function was observed for UMAG_05318 and/or UMAG_11416 (Supplementary Fig. S4). Here, the double mutant KO_UMAG_05318/KO_UMAG_11416 showed reduced virulence on EGB, but not B73. As the single KO strain KO_UMAG_11416 did not show any virulence defect (not shown) and a reduction of virulence for a UMAG_05318 deletion strain had already been reported previously (Schilling et al. 2014), these CSEPs were not further investigated. For all other tested mutants, either no reduction of virulence or a reduction of virulence on all tested maize lines was observed (Supplementary Fig. S4).

To gain more detailed insight into the expression profile of UMAG_02297 during infection progression, relative expression levels were analysed via qRT-PCR on the six different maize lines (Supplementary Fig. S3B). Interestingly, UMAG_02297 was expressed at lowest levels on CML322 throughout the infection process, the maize line on which it was required for full virulence. Hence, high expression levels do not seem to determine the function for virulence. To investigate the relation of expression level of the effector and *U. maydis* virulence, we generated a strain in which UMAG_02297 was expressed under control of the promoter ppit2, which is highly active throughout the infection process and results in a strong overexpression of the gene (Mueller et al. 2013). EGB and CML322 seedlings were infected with Ppit2:UMAG_02297 single and multiple integration strains (Fig. 6D). Interestingly, the overexpression strain showed a maize line-specific virulence defect as well: on CML322, but not on EGB, the multiple integration strain was significantly reduced in virulence compared to strain SG200. This shows that an adjusted expression level of UMAG_02297 is required for virulence on maize line CML322. The finding that neither knockout, nor overexpression of UMAG_02297 had a significant effect on virulence on EGB could suggest that the host targets of this effector might either not be present, or not involved in QDR to *U. maydis* in this maize line.

### Host transcriptional changes induced by UMAG_02297

To investigate which host processes might be influenced by the maize line-specific effector UMAG_02297, leaf samples of CML322 maize seedlings infected with SG200 and KO_UMAG_02297 were analysed by RNA-Seq at 3 dpi. Of all 63477 maize annotated loci, 30637 were expressed in these samples (48%). To assess variability between the samples, we made a multidimensional scaling plot (Fig. 7A). Both *U. maydis*-infected samples formed one cluster highly distinct from the mock-treated samples, indicating that maize gene expression was mostly influenced by infection in general, rather than by the different *U. maydis* strains.

**Figure 7.**
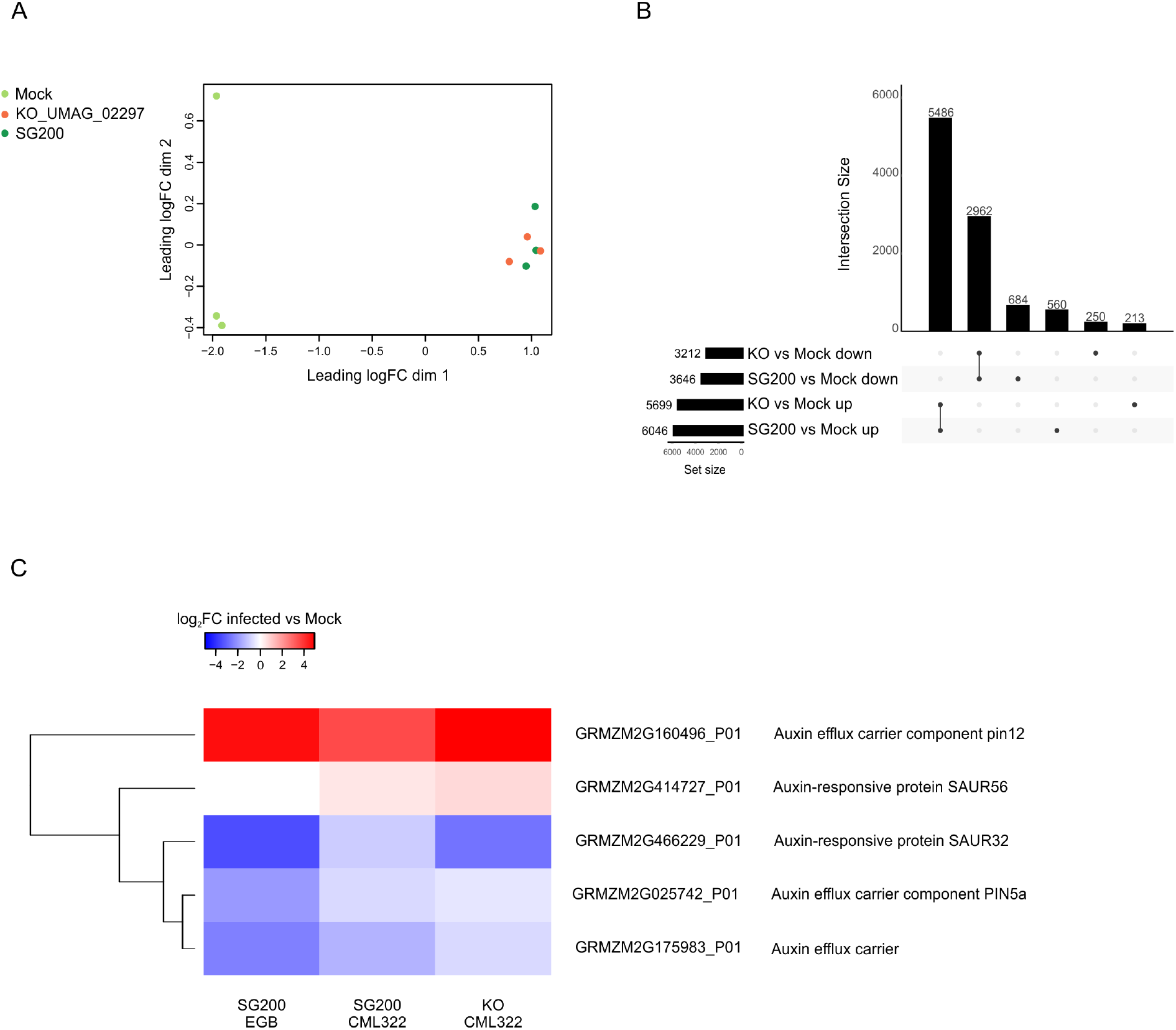
Maize gene expression changes in response to *U. maydis* KO_UMAG_02297. The transcriptome of CML322 maize seedlings infected with SG200, KO_UMAG_02297 and mock was analysed via RNA-Seq 3 days post infection (dpi). KO: knock-out. **A) MDS plot of maize RNA-Seq data.** The top 5000 variable genes were used to calculate pairwise distances between the samples. MDS: multidimensional scaling. **B) UpSet plot of maize genes differentially expressed in response to SG200 and KO_UMAG_02297 infections in comparison to mock.** Genes with a log_2_ expression fold change >0.5 or <-0.5 and adjusted p value <0.05 were considered differentially expressed (DE). In total, 10155 of 30637 expressed genes were DE. Number of differentially expressed genes (DEGs) for each of the 15 possible comparisons is given by set size (horizontal bars). Overlaps of DEGs between comparisons are depicted by connected black dots. Size of overlaps is given by intersection size (vertical bars). **C) Expression profile of auxin-related maize genes in response to *U. maydis* SG200 and KO_UMAG_02297 in EGB and CML322.** Heatmap shows log_2_ expression fold changes of infected vs mock-treated samples.

To identify genes which were uniquely responsive to infection with SG200 or the UMAG_02297 knock-out strain (KO), we compared expression fold changes of the infected samples to the CML322 mock sample (log_2_ expression fold change >0.5, adjusted p value <0.05, Supplementary Data Set S11). Overall, the number of DEGs in response to the KO strain was similar to SG200-infection (SG200: 6046 genes upregulated and 3646 genes downregulated compared to mock;KO strain: 5699 genes upregulated and 3212 genes downregulated compared to mock). Most of the DEGs were jointly regulated: 91% of the upregulated genes (5486) and 81% of the downregulated genes (2962) were equivalently regulated in response to both strains. Only around 9% and 4% (560 and 213 genes) were uniquely upregulated in response to SG200 or KO, respectively, and around 19% and 8% (684 and 250 genes) were uniquely downregulated in response to SG200 or KO, respectively. Taken together, this indicates that maize gene expression is slightly and specifically altered by UMAG_02297. To gain insight into which host processes could be targeted by UMAG_02297, genes uniquely responsive to each of the strains were additionally filtered for genes that were differentially expressed in response to *U. maydis* SG200 infection between CML322 and EGB, where UMAG_02297 was not found to have a function for virulence (426 genes). Within these, genes predicted to encode auxin efflux transporters were strongly enriched (12-fold enrichment, hypergeometric p 0.002, Supplementary Data Set S12). Interestingly, we additionally found several other genes predicted to be related to auxin (Fig. 7C). The auxin-efflux carrier pin12 (GRMZM2G160496_P01) and auxin-responsive SAUR32 (GRMZM2G466229_P01) were similarly regulated in CML322 in response to KO and in EGB in response to SG200, while SAUR56 (GRMZM2G414727_P01) and the auxin-efflux carrier PIN5a (GRMZM2G025742_P01) differed more strongly between the maize lines (SG200- and KO-infected CML322 vs SG200-infected EGB). This observed specific regulation of auxin-related genes identifies the manipulation of the auxin pathway as a potential maize line-specific target of UMAG_02297.

## DISCUSSION

The maize pathogen *U. maydis* serves as a model system to study the molecular mechanisms of biotrophic plant-pathogen interactions and also causes important yield losses in the world’s major crop maize. Unlike in most biotrophic interactions, resistance of maize to *U. maydis* is considered a polygenic, quantitative trait. However, the molecular basis of QDR in the *U. maydis-maize* interaction is mostly unknown. Therefore, the molecular mechanisms underlying QDR in maize and how *U. maydis* interferes with its components were investigated in this study. Plant inoculation experiments revealed that *U. maydis* resistance levels of the NAM founder lines and EGB are highly diverse, which further corroborates the quantitative nature of the *U. maydis-maize* interaction and indicates that several genes are involved in determining resistance or susceptibility. The transcriptome analysis of six *U. maydis*-infected maize lines of different resistance levels offered unprecedented insights into the transcriptional changes associated with host disease resistance.

Resistance levels of the NAM founder lines to other diseases such as Northern corn leaf blight or aphids have been previously analysed, which revealed distinct patterns from the *U. maydis* resistance levels observed in this study. B73 for example is highly susceptible to Northern corn leaf blight, while CML322 is very resistant and Ky21, Oh43 and Tx303 showed medium susceptibility levels (Poland et al. 2011). Aphid resistance was found high on Tx303, Oh43 and Ky21, whereas CML322 was found to be highly susceptible and B73 displayed medium aphid susceptibility (Meihls et al. 2013). For *U. maydis*, CML322 displayed highest resistance levels, followed by B73, and Ky21, Oh43 and Tx303 were moderately to highly susceptible. This suggests that specific rather than general defence mechanisms determine the outcome of maize interactions with different pathogens and pests.

### Maize line-specific gene expression in *U. maydis*

To identify genes involved in QDR in the maize-*U. maydis* interaction, an RNA-Seq analysis of maize lines of highly differing resistance levels colonised by *U. maydis* was performed. This aimed at the identification of fungal genes being expressed in a host-genotype dependent manner, as well as of maize genes being differentially regulated between host genotypes in response to *U. maydis* infection. On the fungal side, the analysis of genes which were differentially expressed between host genotypes showed a significant enrichment of CSEPs. Additionally, CSEPs were significantly enriched in the co-expression module that was negatively correlated to the disease index, i.e. within genes that were upregulated in the more resistant maize lines. Both these findings indicate a predominant role of CSEPs in colonising host lines of different resistance levels and point to an important involvement of CSEPs in targeting components of QDR.

In the co-expression module correlated to colonisation of more resistant host lines, enriched biological processes included mechanisms connected to carbohydrate metabolism, which in plant pathogenic fungi has been directly linked to plant cell wall degradation (Tonukari et al. 2000; Ospina-Giraldo et al. 2003). During *U. maydis* infection, degradation of cell walls was previously found to be important at very early stages to allow initial penetration and intracellular growth, as well as in later stages when plant cell walls need to be loosened to enable cell enlargement for tumour formation, rather than being used as a nutrient source (Doehlemann et al. 2008b; Lanver et al. 2018). Consequently, one could speculate that enhanced cell wall reinforcements or different cell wall compositions might be an additional obstacle the fungus needs to overcome when colonising host lines of higher resistance levels. Several studies have suggested differences in cell wall composition as factors in host-pathogen interactions (Vorwerk et al. 2004; Bacete et al. 2020). *A. thaliana* mutants of the GPI-anchored putative pectate lyase PMR6 were found to be highly resistant to powdery mildew (Vogel et al. 2002). In wheat, variation in pectin composition has been associated with resistance to the stem rust fungus *Puccinia graminis* (Wiethölter et al. 2003). In maize, significant differences in cell wall composition between different lines have been reported (Hazen et al. 2003). A detailed cell wall carbohydrate profiling of all NAM founder lines would allow investigating if more resistant or more susceptible maize lines share similar cell wall compositions and would thereby help to answer the question to which extent natural variation in cell wall composition affects pathogen resistance.

Another functional group of *U. maydis* genes that were correlated to infection of more susceptible lines is linked with ion-transport processes. The exchange of nutrients between cells is predominantly driven by an ion gradient which is produced by the activity of plasma membrane H^+^-ATPases that transport ions through the membrane (Palmgren 1990; Gianinazzi-Pearson et al. 1991; Sondergaard et al. 2004; Wang et al. 2014). During mycorrhizal symbiosis, plant H^+^-ATPases were found to energise nutrient uptake in rice and *Medicago truncatula* (Wang et al. 2014). *U. maydis* contains two H^+^-ATPases that could be involved in nutrient uptake (Robles-Martínez et al. 2013). Together with the previous finding that different nutrient transporters are important virulence factors tied to biotrophic development in *U. maydis* (Wahl et al. 2010; Schuler et al. 2015; Lanver et al. 2018), this could indicate that different availability of nutrients in more resistant vs. more susceptible maize lines is involved in QDR to *U. maydis*.

### Maize processes involved in QDR against *U. maydis*

To gain insight into the host processes which are involved in QDR to *U. maydis*, we analysed genotype-dependent transcriptional changes in response to *U. maydis* in maize lines of distinct resistance levels. The major functional classes within maize DEGs were related to ‘transmembrane transport’ as well as ‘oxidation-reduction’ and ‘protein phosphorylation’. Protein phosphorylation through kinases is a central process for signal transduction in immune responses. Interestingly, kinases have been shown to play important roles in QDR in several cases. Two maize wall-associated kinases, ZmWAK-RLK1 and ZmWAK, confer QDR to northern corn leaf blight and a close relative of *U. maydis, Sporisorium reilianum*,respectively (Hurni et al. 2015; Zuo et al. 2015). Transport processes are essential for plant responses during interactions with pathogens, and several QDR genes encode putative transporters. For example, the ABC transporter encoded by Lr34 confers resistance to diverse fungal pathogens in wheat (Krattinger et al. 2009). Hence, this suggests a possible role for kinases as well as transport processes also in QDR against *U. maydis*. Together, our analysis shows that genes associated to QDR to *U. maydis* involve genes of various functional classes, in line with the complex nature of QDR and the idea that QDR extends beyond pathogen perception (Corwin and Kliebenstein 2017). The finding that only a small fraction of the DEGs was shared with genes previously found to be associated with PTI and ETI in *A. thaliana*, assuming conservation of PTI and ETI between maize and *A. thaliana*,implies that QDR mechanisms are mostly distinct from PTI and ETI gene networks (Dong et al. 2015; Hatsugai et al. 2017; Mine et al. 2018).

Correlation analysis of gene expression to resistance levels via WGCNA again identified processes involved in protein phosphorylation, as well as cell division being upregulated in the more susceptible maize lines in response to *U. maydis*. To build a tumour, the fungus actively triggers cell division and reactivates DNA synthesis in the leaf tissue, which goes along with alterations of genes involved in cell-cycle regulation (Redkar et al. 2015; Matei et al. 2018; Villajuana-Bonequi et al. 2019). In the *A. thaliana-Plasmodiophora brassicae* interaction, which is also accompanied by gall formation, genes involved in cell proliferation were found to be associated with QDR (Jubault et al. 2013). In addition to the obvious involvement in tumour formation, cell-cycle deregulation was also found to have an impact on expression of R genes and can thereby modulate plant defence (Bao et al. 2013). Thus, one could speculate that genes involved in cell division might also play a role in QDR against *U. maydis*.

In the more resistant maize lines, processes involved in photosynthesis were significantly enriched. It was shown previously that *U. maydis* suppresses the induction of photosynthesis-associated genes in infected leaves (Horst et al. 2008; Doehlemann et al. 2008a). This is accompanied by an increase of free hexose levels and a decrease in chlorophyll content, reflecting that the fungus blocks the transition to a photosynthetically active source tissue (Doehlemann et al. 2008a; Matei et al. 2018). Free hexoses within tumour cells are thought to serve as an easily accessible carbon source for the fungus, as well as help to build up osmotic pressure for tumour-cell expansion (Horst et al. 2008; Horst et al. 2010). Infection experiments using maize mutants with distorted starch metabolism showed that alterations in carbon allocation are an important factor influencing *U. maydis* growth and plant defence (Kretschmer et al. 2017). Inefficient suppression of photosynthesis as it is found in the more resistant maize lines might result in changes in carbon allocation as well and consequently lead to a reduced fungal proliferation. However, based on the available data one cannot exclude that the positive correlation of photosynthesis-repression and fungal infection could also be consequence rather than cause of an enhanced susceptibility. Nevertheless, as the developmental stages of the fungus in all our samples were comparable, it is likely that the observed resistance level-specific transcriptional changes directly contribute to the outcome of the quantitative interaction with *U. maydis*.

### Maize line-specific activity of *U. maydis* CSEPs

It has been hypothesized that allelic variation between plant genotypes in genes contributing to resistance or susceptibility likely builds the molecular basis of QDR (Niks et al. 2015). This can lead to altered expression patterns or different modes of defence reactions, but also alter the efficiency an effector can interact with and thereby manipulate its respective host target. Therefore, the targets of pathogen effectors which quantitatively contribute to virulence are potential candidates contributing to QDR and thus, the identification of these targets can help to elucidate the diverse genetic basis of QDR. For one *U. maydis* effector, ApB73, a maize line-specific virulence function has been demonstrated as well, however the differences were only quantitative (Stirnberg and Djamei 2016). One other example strongly supporting the hypothesis that allelic variations in effector targets may be the basis of QDR came from the comparison of the capacity of the EPIC1 effector from two different *Phytophtora* species to suppress their target RCR3, a papain-like cysteine protease. EPIC1 from *P. infestans* was able to inhibit RCR3 from tomato and potato, but not PmEPIC1 from the non-adapted *P. mirabilis*. However, PmEPIC1 was highly effective in inhibiting an RCR3-like protease in *Mirabilis jalapa*. These different specificities resulted from single amino-acid polymorphisms in both the host target and the pathogen effectors, underpinning the importance of ecological effector diversification (Dong et al. 2014).

In this study, we identified a maize line-specific virulence function for the effector gene UMAG_02297, which further substantiates the importance of effectors in targeting components of QDR. The unexpected finding that overexpression of UMAG_02297, similar to its knock-out, resulted in a maize line-specific virulence defect, additionally underlines that manipulation of host processes by effectors requires a fine-tuned adaptation to the host genotype. The analysis of transcriptional changes induced by the UMAG_02297 KO mutant in comparison to wild type infections identified auxin-responsive processes being a possible target of this effector. In general, auxins play a cardinal role in controlling plant growth and development. A role for auxin in the cell enlargement of *U. maydis*-induced tumours has been proposed before, as auxin synthesis as well as auxin-responsive genes are transcriptionally induced during tumour development and auxin levels within *U. maydis*-induced tumours are elevated (Turian and Hamilton 1960; Reineke et al. 2008; Doehlemann et al. 2008a). Additionally, auxin can act as an antagonist of the SA pathway in plant defence, and thereby could promote fungal growth and disease development (Kazan and Manners 2009). Previous studies have identified a large number of auxin-related genes that underlie QDR. For example in the soybean-*Phytophthora sojae* interaction, auxin catabolite accumulation differed between a relatively resistant and a more susceptible soybean cultivar, and the ability of resistant cultivars to cope with auxin accumulation was suggested to play an important role in QDR in this pathosystem (Stasko et al. 2020). In maize, cloning of the causal gene of the Giberella stalk rot resistance QTL *qRfg2* identified ZmAuxRP1, which encodes a plastid stroma-localised auxin-regulated protein, presumably modulating auxin biosynthesis (Ye et al. 2019). Furthermore, increased auxin levels have been generally found to lead to enhanced susceptibility to several biotrophic pathogens (Navarro et al. 2006; Wang et al. 2007; Mutka et al. 2013) and a few pathogen effectors that target auxin-related processes have been identified so far. The *P. syringae* effector AvrRpt2 for example initiates auxin signalling through degradation of auxin/indole acetic acid (IAA) proteins (Cui et al. 2013), and the effector PSE1 from *Phytophthora parasitica* modulates local auxin levels through altered distribution of auxin efflux transporters (Evangelisti et al. 2013). Together, this renders auxin-related processes an interesting and promising possible target of UMAG_02297, which will be tested in future studies.

Even though fundamental progress in the field of plant-pathogen interactions has been achieved through reductionist approaches, the need for more holistic studies that integrate the natural variation in both pathogen and host has become increasingly evident. As a first step, the present study revealed the influence of different host genotypes on *U. maydis* virulence and thereby found that activity and function of effector genes are specifically dependent on the host line. For future studies it will be seminal to investigate the intraspecific variability of the maize line-specific effector genes in *U. maydis:* are different variants of the same effectors functional in different host genotypes and how does a highly specific, biotrophic pathogen like *U. maydis* co-adapt with its host maize in different ecologic background.

## MATERIALS AND METHODS

### Plant growth conditions, fungal infections and collection of samples

*Zea mays* L. lines Early Golden Bantam (Olds Seeds, Madison, WI, USA) and the inbred founder lines of the Nested Association Mapping (NAM) population (Yu et al. 2008; McMullen et al. 2009; North Central Regional Plant Introduction Station, IA, USA) were used for infections.

Virulence assays of *U. maydis* on *Z. mays* were performed as described in Redkar and Doehlemann (2016) in three independent biological replicates. Virulence symptoms were scored 12 dpi using the disease rating scheme developed previously (Kämper et al. 2006). Disease indices were calculated as follows: The number of plants sorted into categories ‘small tumours’, ‘normal tumours’, ‘heavy tumours’ and ‘dead plants’ were multiplied by the number of the category (1; 3; 5; 7, respectively), summed and then divided by total number of infected plants: {[(symptom category X × number of plants in category X)X+(…)Y-Z]/total number of plants}. The resulting value for SG200 was set to unity. Indices of deletion mutants are given relative to the score for SG200. An unpaired t-test was used to calculate the statistical significance of the differences in disease indices between mutant strains and SG200.

Samples of 20 infected maize seedlings were collected at 1, 3, 6 and 9 dpi in three independent biological replicates. For 1 dpi, 2 cm sections of the third leaves were excised 0.5 cm below the infection holes. At later time points (3, 6 and 9 dpi), 5 cm sections of the third leaves were excised 1 cm below the infection holes. Comparable sections were harvested for mock-treated controls. For each sample, leaf sections of 20 different plants were pooled, immediately frozen in liquid nitrogen and stored at −80 °C.

### Generation of fungal strains

All *U. maydis* strains used in this study were derived from the solopathogenic strain SG200 (Kämper et al. 2006). To generate *U. maydis* knock-out mutants, the CRISPR-Cas9 system using the non-integrative, self-replicating backbone plasmid pMS73 was employed (Schuster et al. 2016; Schuster et al. 2018). All target sequences for the guide RNA constructs were designed using the E-CRISP tool (www.e-crisp.org; Heigwer et al. 2014) with medium stringency settings. For integration into the *ip* locus of *U. maydis*, plasmids derived from p123 were used and linearised within the cbx gene before transformation into the respective *U. maydis* strains. Transformation of *U. maydis* was carried out as described previously (Schulz et al. 1990). To identify strains with Cas9-induced mutations leading to gene knock-out, the respective loci were amplified and sequenced with gene-specific primers. The stable integration of p123-based plasmids into the *ip* locus was verified by Southern blot analysis. All complementation constructs were integrated in single copy into the *U. maydis ip* locus.

### Microscopic analyses

To visualise *U. maydis* infection progression, infected maize leaf samples were stained with WGA-AF488 and propidium iodide as described in Redkar et al. (2018) and analysed using a Zeiss Axio Zoom V16 using the GFP filter for WGA-AF488 and the DsRed filter for propidium iodide visualization. Image processing was done using ImageJ.

### Extraction of nucleic acids and RNA sequencing

For the isolation of genomic DNA from *U. maydis*, a phenol-based extraction method was used (Hoffman and Winston 1987). For isolation of nucleic acids from maize leaves, pooled leaf sections of the individual maize samples were homogenised using a mortar and pestle under constant liquid N_2_ cooling. Isolation of genomic DNA from leaf powder was performed using the MasterPure™ Complete DNA and RNA Purification Kit from Epicentre (Epicentre, Chicago, USA) according to manufacturer’s instructions. For isolation of total RNA, TRIzol^®^ reagent (Invitrogen, Darmstadt, Germany) was used according to the manufacturer’s instructions. To approximately 400 μl of homogenised tissue 1 ml TRIzol^®^ reagent was added immediately. To eliminate genomic DNA contamination, the Turbo DNA-Free™ Kit from Ambion (Ambion Life technologies™, Carlsbad, USA) was used according to the manufacturer’s instructions. Sequencing library preparation was done using the Illumina TruSeq mRNA stranded Kit (Illumina, San Diego, USA) or NEB Next^®^ Ultra™ RNA Library Prep Kit (NEB, Ipswich, USA). Illumina sequencing of mRNA was performed with 150 bp paired-end reads at the Cologne Center for Genomics (CCG, Cologne, Germany) on an Illumina HiSeq 4000 (Illumina, San Diego, USA) and at Novogene (Peking, China) on an Illumina NovaSeq 6000 (Illumina, San Diego, USA).

### RNA-Seq data analysis

By Illumina sequencing of mRNA libraries, approximately 60 million 150 bp paired-end reads per *U. maydis*-infected sample and 40 million paired-end reads per mock-treated sample were created. The reads were filtered using the Trinity software (v2.9.1) option trimmomatic with standard settings (Grabherr et al. 2013). The reads were then mapped to the reference genome using Bowtie 2 (v2.3.5.1) with the first 15 nucleotides on the 5’-end of the reads being trimmed (Langmead and Salzberg 2012). As reference genome the *U. maydis* genome assembly (Kämper et al. 2006) and the *Z. mays* B73 version 3 (Schnable et al. 2009) genome assembly were used simultaneously. Reads were counted for *U. maydis* and *Z. mays* loci using the R (www.r-project.org) package Rsubread (v1.34.7) (Liao et al. 2019). On average, 640000 mapping read counts were for the *U. maydis* genome were found per sample in the dataset of different maize lines (1.3% of total read counts) and 783000 read counts for the data set of CML322 infected by SG200 or KO_UMAG_02297 (1.8% of total read counts). For maize, approximately 50 million read were counted for the *U. maydis*-infected samples and 43 million for the mock samples. Pre-filtering was applied to keep only genes with at least 10 counts in 3 samples (6284 genes for *U. maydis*, 40056 genes for the data set of different maize lines and 30637 genes for the data set of CML322 infected by SG200 or KO_UMAG_02297). Counts for *U. maydis* or maize were normalised and differential gene expression was analysed by DESeq2 v1.26.0 (differential expression analysis for sequence count data 2, Love et al. 2014) in R. For *U. maydis*, the design formula was `~genotype`, for maize, the design formula was `~genotype + condition` + genotype:condition to identify differences in condition effects (SG200-infected vs mock) between genotypes. Genes with a log_2_ expression fold change >0.5 and Benjamini-Hochberg-adjusted p value <0.05 were considered differentially expressed. To identify co-expressed genes, a weighted gene co-expression network analysis (WGCNA) was done using the WGCNA package (v1.69) (Zhang and Horvath 2005; Langfelder and Horvath 2008) in R. Only genes with at least 10 counts in 50% of the analysed maize samples or in 90% of the analysed *U. maydis* samples were considered. For *U. maydis*, 4013 genes and for maize 29729 genes passed this filtering. Log_2_-transformed DESeq2-normalised counts were used as input for the network analysis. The function blockwiseModules was used to create a signed network of a Pearson-correlated matrix; only positive correlations were considered. For *U. maydis*, all genes were treated in a single block. For maize, the maximum blocksize was set to 15000. The soft power threshold was set to 4 for *U. maydis* and for maize because this was the lowest power needed to reach scale-free topology (R^2^= 0.901 and 0.871 respectively). Modules were detected using default settings with a mergeCutHeight of 0.15 and a minimal module size of 25 genes. For each module, the expression profile of the module eigengene was calculated, which represents the modules by summary expression profiles of all genes of a given module. For each gene and module eigengene, the Pearson correlation to the disease index of the different maize lines was calculated (= gene significance for the trait).

### GO enrichment analysis

GO term enrichment analysis (Ashburner et al. 2000; The Gene Ontology Consortium 2017) for *U. maydis* was performed with the Gene Ontology Panther Classification System (Mi et al. 2019) using a p value cut-off of <0.05. For the enrichment analysis of the modules correlated to the disease index, only genes were considered that had a gene significance for disease index > 0.5 or < −0.5 and p value <0.05. For maize, Gene Ontology (GO) terms were annotated to the version 3 protein annotation of maize line B73 using InterProScan (v5.42-78.0) (Schnable et al. 2009; Jones et al. 2014; El-Gebali et al. 2019). Significance of GO term enrichments in a subset of genes were calculated for all expressed genes with a Fisher’s exact test with the alternative hypothesis being one-sided (greater).

### Quantitative reverse-transcriptase PCR (qRT-PCR) and quantitative PCR (qPCR)

The qRT-PCRs/qPCR reactions were set up using the GoTaq^®^ qPCR Mastermix (Promega, Madison, Wisconsin, USA) according to the manufacturer’s instructions in a total volume of 15 μl. All qRT-PCRs/qPCRs were performed in an iCycler system (Bio-Rad, Munich, Germany) with the following program: 95 °C / 2 min-(95 °C / 30 s-62 °C / 30 s-72 °C / 30 s) x 45 cycles. For gene expression analysis by qRT-PCR, cDNA was synthesised from 1-5 μg of template RNA using he Thermo Scientific RevertAid H Minus First Strand cDNA Synthesis Kit by Thermo Fisher Scientific (Waltham, Massachusetts, USA) according to the manufacturer’s instructions. *U. maydis* ppi was used as the normalization gene and relative expression values were calculated using the 2^−ΔCt^ method. For fungal biomass quantification, DNA extracted from infected maize leaves was subjected to qPCR analysis with maize-specific GAPDH and *U. maydis*-specific ppi primers. The relative fungal biomass was calculated as the ratios of *U. maydis* DNA to maize DNA (2^−ΔCt^).

### Mapping of maize genes to Arabidopsis

For comparison to genes previously described to be involved in PTI or ETI in Arabidopsis, mapping of maize gene IDs to Arabidopsis was performed on the Monocots PLAZA 4.0 workbench (https://bioinformatics.psb.ugent.be/plaza/, Van Bel et al. 2018) using the PLAZA orthologous genes integrative method with standard settings and a minimum number of required evidence types of three.

## Supporting information

Supplementary data sets

## Data availability

RNA sequencing data has been submitted to NCBI Genbank and are available under the following links: httpsXXXXXXXXX

## ACKNOWLEDGEMENTS

We are grateful to Benjamin Stich for generously supplying Ky21 and Tx303 seeds, to Elaine Jaeger for helping with the initial acquirement of NAM founder lines and Mariana Schuster for providing *U. maydis* strains. We thank Henriette Läβle for excellent help with the project. Selma Schurack received support by the International Max Planck Research School (IMPRS) of the MPIPZ, Cologne, Germany. This work was funded by the European Research Council under the European Union’s Horizon 2020 research and innovation program (consolidator grant conVIRgens, ID 771035), as well as funding by the Deutsche Forschungsgemeinschaft (DFG, German Research Foundation) under Germany’s Excellence Strategy-EXC-2048/1-Project ID: 390686111. Jasper Depotter is supported by the Research Fellowship Programme for Postdoctoral Researchers of the Alexander von Humboldt Foundation. Marco Thines and Deepak K. Gupta are supported by the LOEWE initiative of the government of Hesse, in the framework of the centre for Translational Biodiversity Genomics (TBG).

## SUPPLEMENTARY CAPTIONS

**Figure S1. WGA-AF488/propidium iodide co-stained maize leaves infected with *U. maydis*.** Samples were collected at 1, 3, 6 and 9 days post infection (dpi). Fungal hyphae were visualised by staining with WGA-AF488 (green), plant cell walls were visualised by staining with propidium iodide (red). Scale bars = 200 μm.

**Figure S2. Identification of genes previously associated with PTI or ETI immune responses within maize DEGs.** *A. thaliana* orthologues of maize differentially expressed genes (DEGs) were examined for overlaps to genes previously identified to be associated with PAMP-triggered- (PTI, left) or effector-triggered immunity (ETI, right) (Dong et al. 2015; Hatsugai et al. 2017; Mine et al. 2018).

**Figure S3. Expression of UMAG_02297 during disease progression in different maize lines.** Quantification of UMAG_02297 relative expression via qRT-PCR during infection progression at 1, 3, 6, and 9 days post infection (dpi). Solid points give mean ratios of UMAG_02297 to ppi (2^−ΔCt^) of three biological replicates. Transparent points give individual values; error bars denote the standard deviation.

**Figure S4. Virulence functions of candidate maize line-specific effectors.** Double and quadruple knock-out (KO) mutant strains of selected maize line-specific effectors were injected into maize seedlings of the indicated line and symptoms were scored 12 days post infection (dpi). Gene names are given at the top. KO refers to the respective CRISPR/Cas9 knock-out strain. Gene names separated by slash indicate double or quadruple KO of these genes. KO/C indicates that a single copy of the respective genes was introduced into the KO strain for complementation. Disease indices reflect disease symptom severity and are shown in relation to SG200, which was set to unity. Asterisks label significant reduction in disease index compared to SG200 (student’s t-test, p<0.05). All experiments were performed in three independent biological replicates. Average number of infected plants per strain and maize line: 90. Strain KO_UMAG_11444/KO_UMAG_11002 was kindly provided by Mariana Schuster.

**Supplementary Data:** Supplementary data sets S1-S12

