## Supplementary Figures for "Transcriptome analysis in the maize-*Ustilago maydis* interaction identifies maize-line-specific activity of fungal effectors"

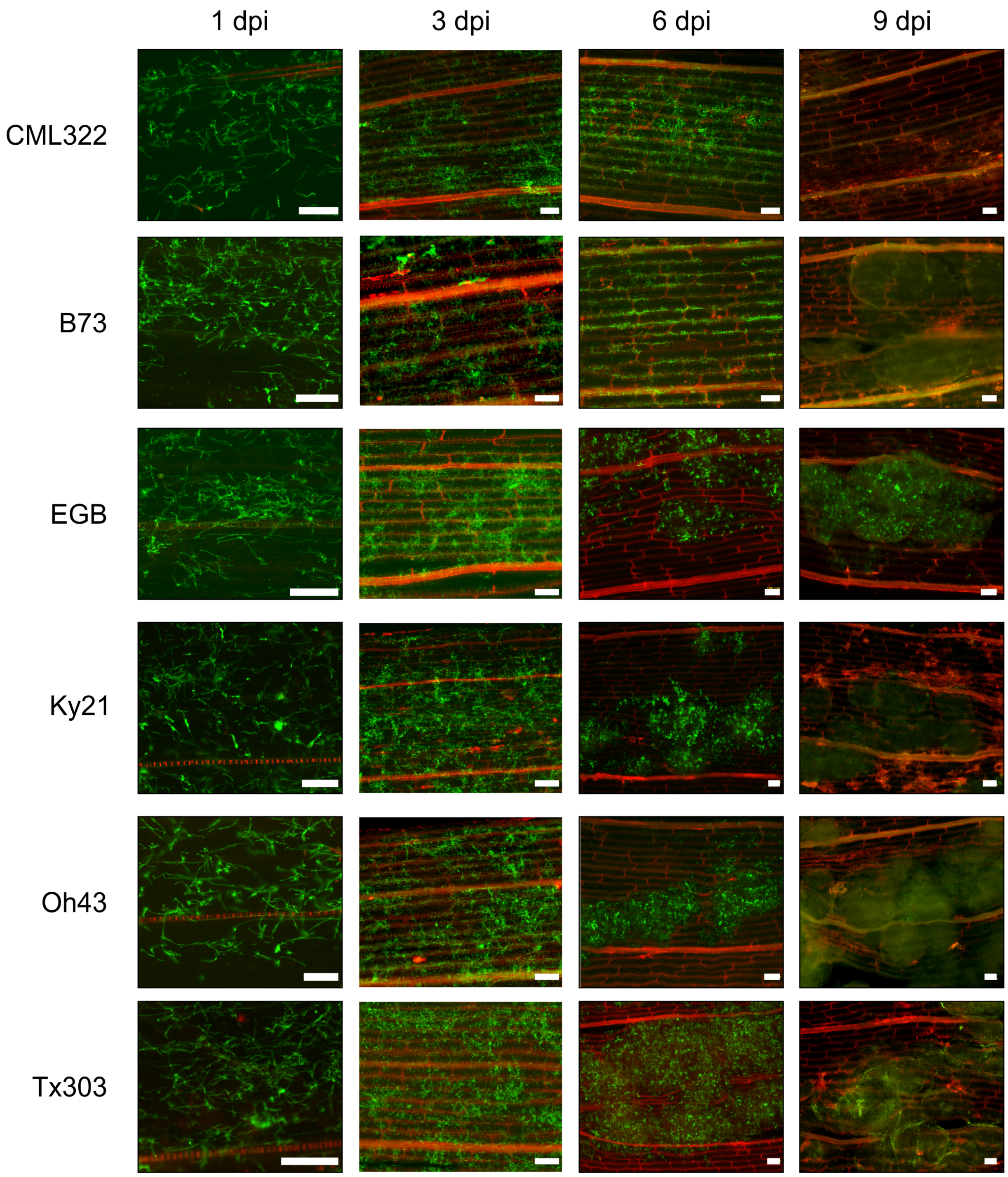


**Figure S1**. **WGA-AF488/propidium iodide co-stained maize leaves infected with *U. maydis*.** Samples were collected at 1, 3, 6 and 9 days post infection (dpi). Fungal hyphae were visualised by staining with WGA-AF488 (green), plant cell walls were visualised by staining with propidium iodide (red). Scale bars = 200 µm.

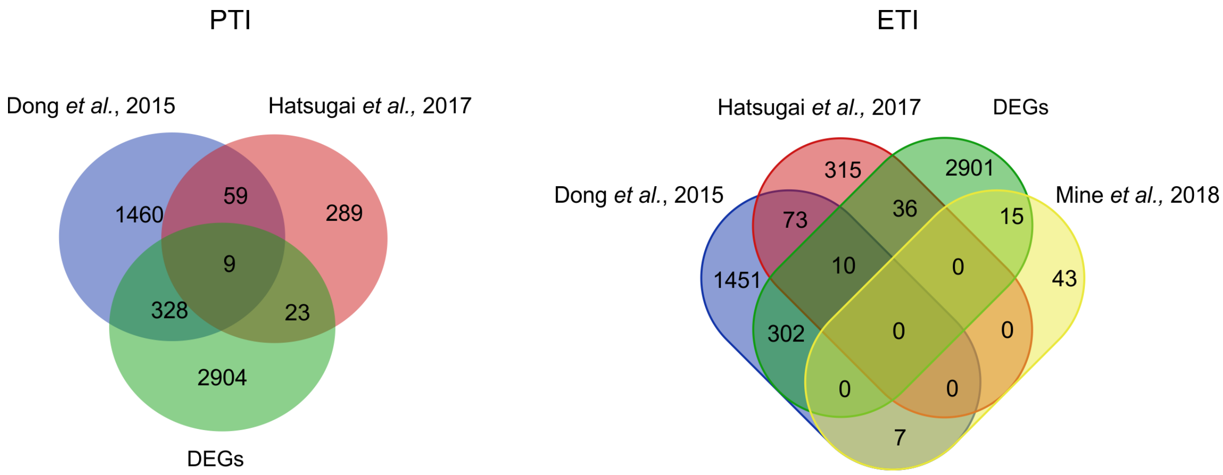


**Figure S2. Identification of genes previously associated with PTI or ETI immune responses within maize DEGs.** *A. thaliana* orthologues of maize differentially expressed genes (DEGs) were examined for overlaps to genes previously identified to be associated with PAMP-triggered- (PTI, left) or effector-triggered immunity (ETI, right) (Dong et al. 2015; Hatsugai et al. 2017; Mine et al. 2018).

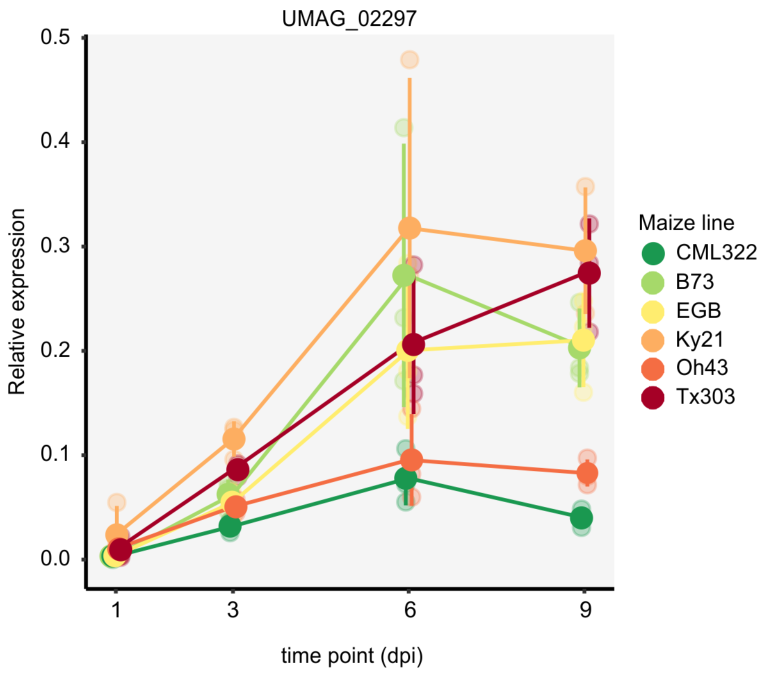


**Figure S3. Expression of UMAG_02297 during disease progression in different maize lines.** Quantification of UMAG_02297 relative expression via qRT-PCR during infection progression at 1, 3, 6, and 9 days post infection (dpi). Solid points give mean ratios of UMAG_02297 to ppi ($2^{-\Delta Ct}$) of three biological replicates. Transparent points give individual values; error bars denote the standard deviation.

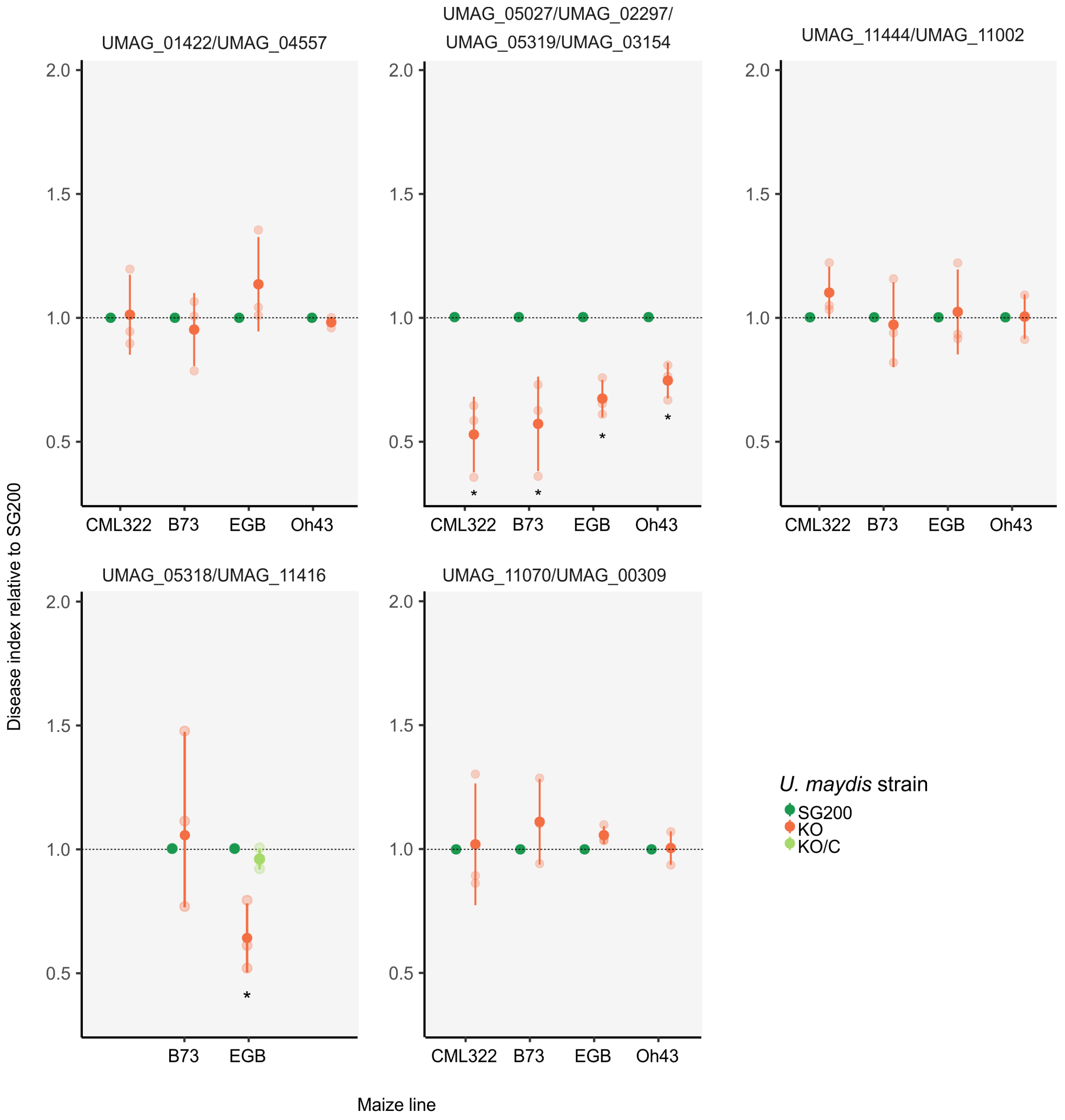


**Figure S4. Virulence functions of candidate maize line-specific effectors.** Double and quadruple knock-out (KO) mutant strains of selected maize line-specific effectors were injected into maize seedlings of the indicated line and symptoms were scored 12 days post infection (dpi). Gene names are given at the top. KO refers to the respective CRISPR/Cas9 knock-out strain. Gene names separated by slash indicate double or quadruple KO of these genes. KO/C indicates that a single copy of the respective genes was introduced into the KO strain for complementation. Disease indices reflect disease symptom severity and are shown in relation to SG200, which was set to unity. Asterisks label significant reduction in disease index compared to SG200 (student's t-test, $p<0.05$). All experiments were performed in three independent biological replicates. Average number of infected plants per strain and maize line: 90. Strain KO_UMAG_11444/KO_UMAG_11002 was kindly provided by Mariana Schuster.
